# Oral administration of 1,10-phenanthroline-5-amine (PAA) significantly reduced amyloid plaque burden compared with untreated APP/PS1 mice

**DOI:** 10.64898/2026.07.27.740397

**Authors:** Larry Schmued, Bryan Maloney, Calvert Schmued, Kaitlyn Gregerson, Debomoy K. Lahiri

**Affiliations:** Histo-Chem Inc, White Hall, AR, USA; Departments of Psychiatry; Departments of Medical & Molecular Genetics, Stark Neurosciences Research Institute, Indiana University School of Medicine, Indian Alzheimer’s Disease Research Center, Indianapolis, IN, USA

**Keywords:** Aging, APP/PS1 mice, amyloid load, brain, gangliosides, metal chelators, neurodegeneration

## Abstract

**Background:** We previously demonstrated that 1,10-phenanthroline-5-amine (PAA) significantly reduced the number and size of amyloid plaques in one-year-old APP/TAU mice. The primary objective of the present study was to validate these findings in the APP/PS1 mouse model using a larger cohort of animals. A second objective was to determine whether PAA binds directly to amyloid plaques in brain tissue sections.

**Methods:** For the *in vivo* studies, APP/PS1 mice received daily oral PAA or vehicle treatment and were euthanized at one year of age. Brains were collected, fixed, cryosectioned, and stained with hydroxyquinoline oxalate (HQ-O) to visualize amyloid plaques. For the *in vitro* studies, brain tissue sections were incubated in a PAA solution. Double labeling with PAA and HQ-O was performed on the same tissue sections to compare plaque labeling patterns.

**Results:** Daily oral administration of PAA produced a significant reduction in both the number and size of amyloid plaques compared with untreated control mice. In vitro incubation of tissue sections with PAA resulted in red fluorescent labeling of all amyloid plaques. Double-labeling studies showed that PAA labeled plaques are more extensive than HQ-O in frozen tissue sections, whereas no such difference was observed in paraffin-embedded sections.

**Conclusions:** These findings extend our previous observation that chronic oral administration of PAA significantly reduces amyloid plaque burden in vivo. In addition, the in vitro studies demonstrate that PAA binds directly to amyloid plaques. The mechanism of PAA binding may involve interactions with transition metals incorporated within amyloid plaques and/or the sialic acid moieties of plaque-associated gangliosides.

## Background

Alzheimer’s disease (AD) is the most common cause of age-associated dementia (1). AD is a major unsolved problem with limited treatment options. The number of people living with AD and related dementias (ADRD) worldwide was estimated at 47 million in 2015, with an estimated prevalence reaching 132 million in 2050 (2). In the US alone, over 6 million individuals live with AD, and numbers are expected to rise with increased life expectancy. However, the biochemical cascade of events leading to AD remains elusive. Our goal is to identify mechanisms in AD, which may be common to ADRD, and devise novel preventive, therapeutic and diagnostic strategies for AD.

AD is characterized by amyloid plaques, neurofibrillary tangles, inflammation, and ultimately neuronal death. The neurofibrillary tangles are composed of hyperphosphorylated Tau filaments, and the amyloid plaques contain Aβ aggregates and a variety of proteins, lipids, sugars, and metals. Several metals may contribute to the emergence of AD, including lead (3–6), iron (7–11), copper (12–16), manganese (17–19), zinc (11, 14, 20–23), and lithium (24–29). Of these, lithium, a monovalent metallic ion, may be protective, while the rest, divalent or oligo-valent, appear to pose hazards. Various strategies have been employed in an effort to slow, stop or reverse the amyloidosis process. Although some compounds have shown promise in mouse models of AD, thus far nothing has translated to effective human therapy. Several cellular and molecular targets enrich our understanding of mechanisms leading to AD (30–33). However, no effective exogenous treatment has yet been discovered or engineered, despite promising initial data (34–38).

Metals play a critical, though complex, role in the neurodegeneration that leads to AD (17, 39, 40). While many of these same metals are essential for healthy brain function, imbalances or disruptions in metal homeostasis may contribute to the development and progression of AD. Specifically, certain metals can impact the formation of amyloid plaques and neurofibrillary tangles, hallmarks of the disease, and impact on neuronal health and function. Studies on amyloid aggregation typically look for a molecular ‘seed’ that facilitates this aggregation. One strong candidate class is certain transition metals including Fe, Zn, Cu, and Mn (41, 42). Such metals can accelerate fibrillization of Aβ *in vitro* (17, 43) and are found at elevated concentrations within amyloid plaques (14, 42, 44–48) in humans and transgenic mice. A variety of metal chelators showed promise for attenuating amyloid lesions in transgenic mice (48–58), but few were tested in human subjects. Nevertheless, several chelators were promising in reducing amyloidosis in clinical studies, the first being deferoxamine (59). Another metal chelator reported to reduce amyloid plaques in humans is clioquinol (60, 61) and its analogues VK-28, HLA-20, PBT2 and M-30. One limitation to the clinical use of clioquinol is its tendency to cause subacute myelopathic neuropathy (SMON) in humans (61). While deferoxamine is typically administered intravenously or intramuscularly(59), intranasal administration may be practical (59).

Other molecules whose dysregulation were implicated in AD include certain glycosylated and sialylated molecules. About 1/3 of the glycosylated molecules in the brain are glycoproteins while the remaining 2/3 are glycolipids. The most abundant glycolipid is monosialotetrahexosylganglioside (GM1), which is heavily glycosylated and sialylated. As a major component of neural plasma membranes, GM1 supports neuronal growth, differentiation, and protection, and it can cross the blood-brain barrier. Aβ can bind to gangliosides to form GAβs, which could result in Aβ oligomers forming amyloid aggregates (62, 63). Numerous subsequent studies support this hypothesis (64, 65). Circular dichroism indicated that Aβ specifically binds to ganglioside-containing membranes, which induces an initial unordered configuration transitioning to a β-sheet configuration at higher ganglioside concentrations (66, 67). Ramin spectroscopy suggested that Aβ is first bound to a sialic acid group of GM1 (68). The ratio of Aβ to GM1 on raftlike lipid bilayers composed of GM1, cholesterol and sphingomyelin has been investigated using circular dichroism. Such studies show that at Aβ:GM1 ratios of less than 0.013 only the helical form of Aβ exists. However, at ratios between approximately 0.013 and 0.044 both the helical and β-sheet configuration exist, while ratios above 0.044 produce a form of Aβ in the pleated sheet configuration that will form larger aggregates (67, 69). Although the specifics of how Aβ binds to GM1 is not fully known, the negatively charged sialic acid in GM1 binds to positively charged lysine in the Aβ monomer (70).

Our previous study (71) showed that the all-cationic metal chelator PAA will reduce amyloid plaque burden by approximately 1/3 in an APP/Tau mouse model of AD. The compound was chosen for study because of its unusual chemical properties. Fig. 1 illustrates the compound’s structure showing the two tertiary nitrogen groups in the aromatic portion of the molecule, which serve to chelate a wide variety of oligo-valent metals, and the primary nitrogen of the amino group, which makes the molecule a relatively strong cation. In this respect, PAA differs from other metal chelators previously studied as possible AD therapeutics. The present study complements the original study. One difference was the use of APP/PS1 mice, since the APP/Tau model used in the earlier study produced fewer plaques. In addition, this study also involves *in vitro* studies on the histochemical binding and localization of PAA to amyloid plaques in fixed brain tissue sections. It must be noted that the aggressive temperament of APP/Tau mice resulted in the deaths of 3 littermates prior to dosing, resulting in fewer animals than used in the present study.

**Fig. 1.**
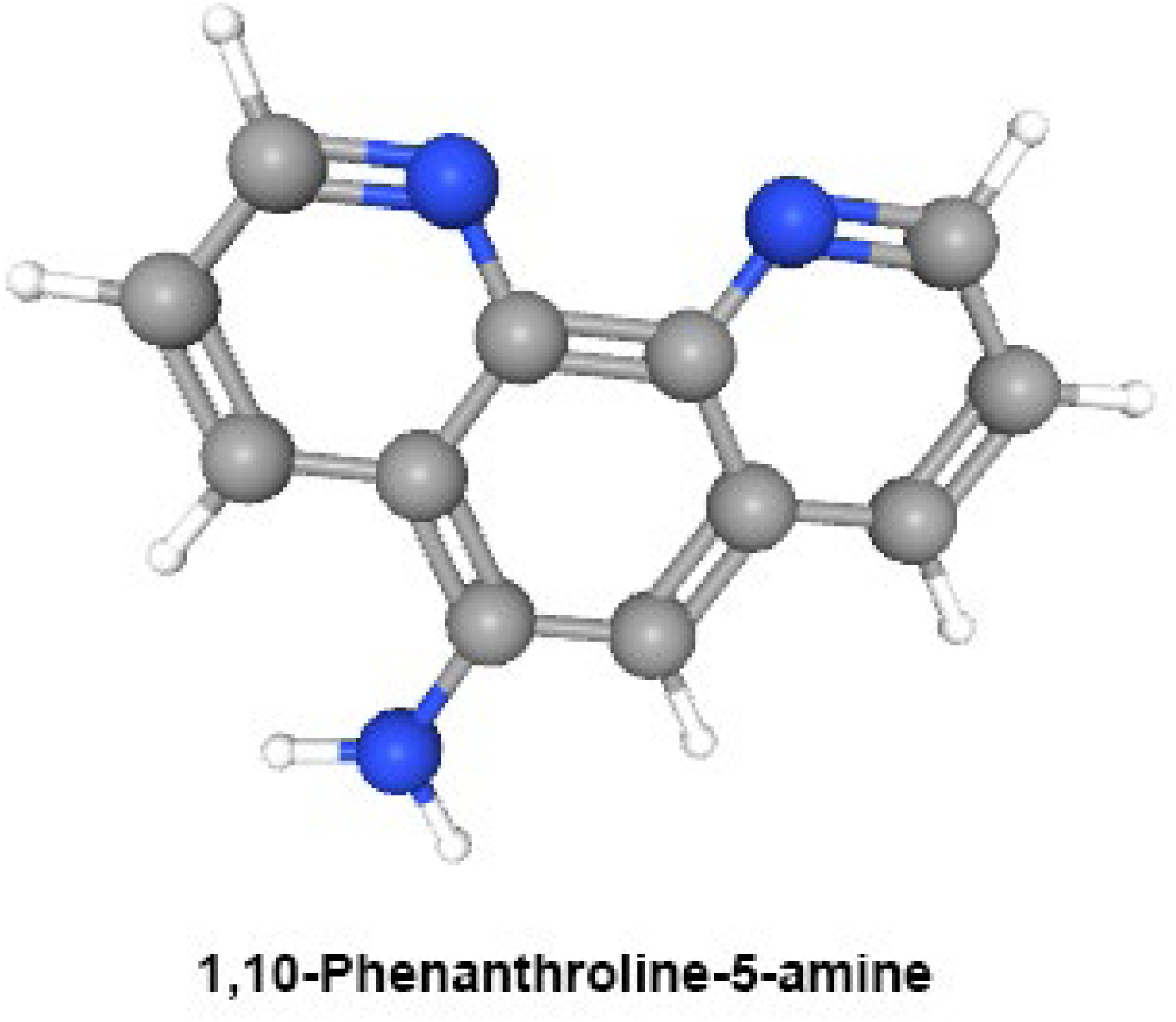
Chemical structure of PAA. Primary and tertiary nitrogen atoms are indicated in blue.

Histological localization of amyloid plaques can be accomplished with a number of different tracers. The majority of these exhibits an affinity for Aβ aggregates in a pleated sheet configuration. Such tracers include Congo Red (72), Thioflavin S (73), Amylo-Glo (74), FSB (75), and antibodies to Aβ (44). But less common are probes that show an affinity for other plaque epitopes. This includes those that bind to n-acetyl neuraminic acid containing proteins and lipids, such as the periodic acid-Schiff’s reagent (PAS) (76), and Ruthenium Red (77) staining techniques. The Fluoro-Jade dyes also label a component of amyloid plaques (78) that likely corresponds to degenerating neurites. Also, plaques can be labeled with certain metal chelators such as 8-hydroxyquinoline oxalate (HQ-O) (44). In addition to demonstrating a reduction of plaque burden by PAA in transgenic mouse models of AD, this study represents the first case of PAA being used as a histological probe for localizing amyloid plaques.

## Methods

### Animals

The *in vivo* portion of the present study used the same experimental design as the original study (71) but differed only in the larger number of animals used and in the use of the double transgenic female APP/PS1 C57BL/6NTac.CBA-Tg(Thy1-PSEN1*M146V,-APP*Swe)10Arte mouse model of AD (Taconic, Rensselaer NY) which develop brain lesions composed of amyloid plaques typical of AD. The animals were used in accordance with animal use protocol HC-AD-029-B and were overseen by the Histo-Chem Institutional Animal Care and Use Committee. Animals were individually housed and put on a 12 hr on / 12 hr off light cycle. The animals had full time access to standard diet and were given a weekly supplement of peanuts to optimize nutrition and vitality.

The mice were allotted into either an untreated control group or a group orally dosed with PAA solution from 4 months of age until their sacrifice at 12 months of age. PAA (Sigma-Aldrich Co., St Louis, MO) was administered via drinking water: 610 mg PAA was added to 1L of water plus 1.2ml of 25% HCl to solubilize the compound. Treated animals were given free access to the dosed drinking water. No difference was seen between the control and treated mice regarding average weight, food or water intake. The PAA dosing solution had a clear amber color, a slightly tart taste and a pH of 4.5. Both groups of animals consumed an average of 3.25 ml of water or dosing solution daily. This resulted in the experimental group receiving an average total oral daily dose of 3.9 mg per animal, which translates into an average daily oral dose of 97 mg/kg of PAA.

### Histology

At 1 year of age all animals were euthanized and perfused vascularly with 10% formalin in PBS. Their brains were removed and placed in the same fixative solution plus 20% sucrose. The brains were cut on a Leica CM 3050 S cryostat into 25μm thick sections through the entire cortex. Tissue sections were then mounted on gelatin coated slides and air dried. A subset of these sections was then delipidized by immersion in solutions of xylene followed by ethanol for 10 minutes each. The following histochemical stains were performed:

#### HQ-O

One out of every 4 tissue sections was mounted onto gelatin coated slides and then stained with HQ-O, which involved incubating slides in a .025% solution of HQ-O (Histo-Chem Inc., Jefferson AR) for either 90 min at 55°C or 5 hours at room temperature (44). Slides were rinsed in distilled water, dehydrated on a slide warmer, cleared for about 1 min in xylene and coverslipped with DPX mounting media. The sections were examined under blue light excitation. Plaque images were captured and their numbers and areas determined.

#### PAA

This histological method involved first incubating slide mounted tissue sections in a .025% solution of PAA in .02M acetate buffer (pH 4.5) at room temperature for 90 minutes. Sections were rinsed through two 3 min changes of distilled water and were then air-dehydrated, cleared in xylene and coverslipped with DPX mounting media. Sections were examined under green, blue and UV illumination. Following PAA staining, a subset of slides was immersed in a .1% solution of EuCl_3_ in distilled water for 30 min at 55°C. These slides were then rinsed in distilled water, air-dried, coverslipped and examined under UV illumination.

### Multiple labeling

To help characterize PAA plaque labeling, the stain was combined simultaneously with HQ-O by dissolving 16 mg of PAA and 25 mg of HQ-O in 100 ml of distilled water. Tissue sections were immersed in this mixture for 5 hours at room temperature, rinsed in distilled water, air-dried, xylene cleared and coverslipped with DPX mounting media.

### Statistical Analysis

Total area, plaque counts, and mean area per plaque were modeled by generalized mixed-level linear models with Group (Control vs. Experimental) as the fixed effect. To account for technical replication and necessary segmentation of the study, random slopes and intercepts were clustered by mouse and by segmentation, section, and slide. Since they were count data, plaques were modeled under the generalized Poisson distribution (79). All models were tested by ANOVA and estimations made of marginal means and Cohen’s *d* effect sizes. Modeling and statistics were performed with the R environment, and the glmmTMB (80), MuMIn (81), and emmeans (82) packages.

## Results

### Oral administration of PAA significantly reduced amyloid plaque burden in vivo

PAA treatment significantly reduced amyloid plaque burden by decreasing plaque seeding, plaque size, and total plaque area. All animals survived until the planned sacrifice at one year of age. Visual inspection of HQ-O-stained brain sections revealed a lower density of amyloid plaques in PAA-treated mice compared with untreated controls (Fig. 2).

**Fig. 2.**
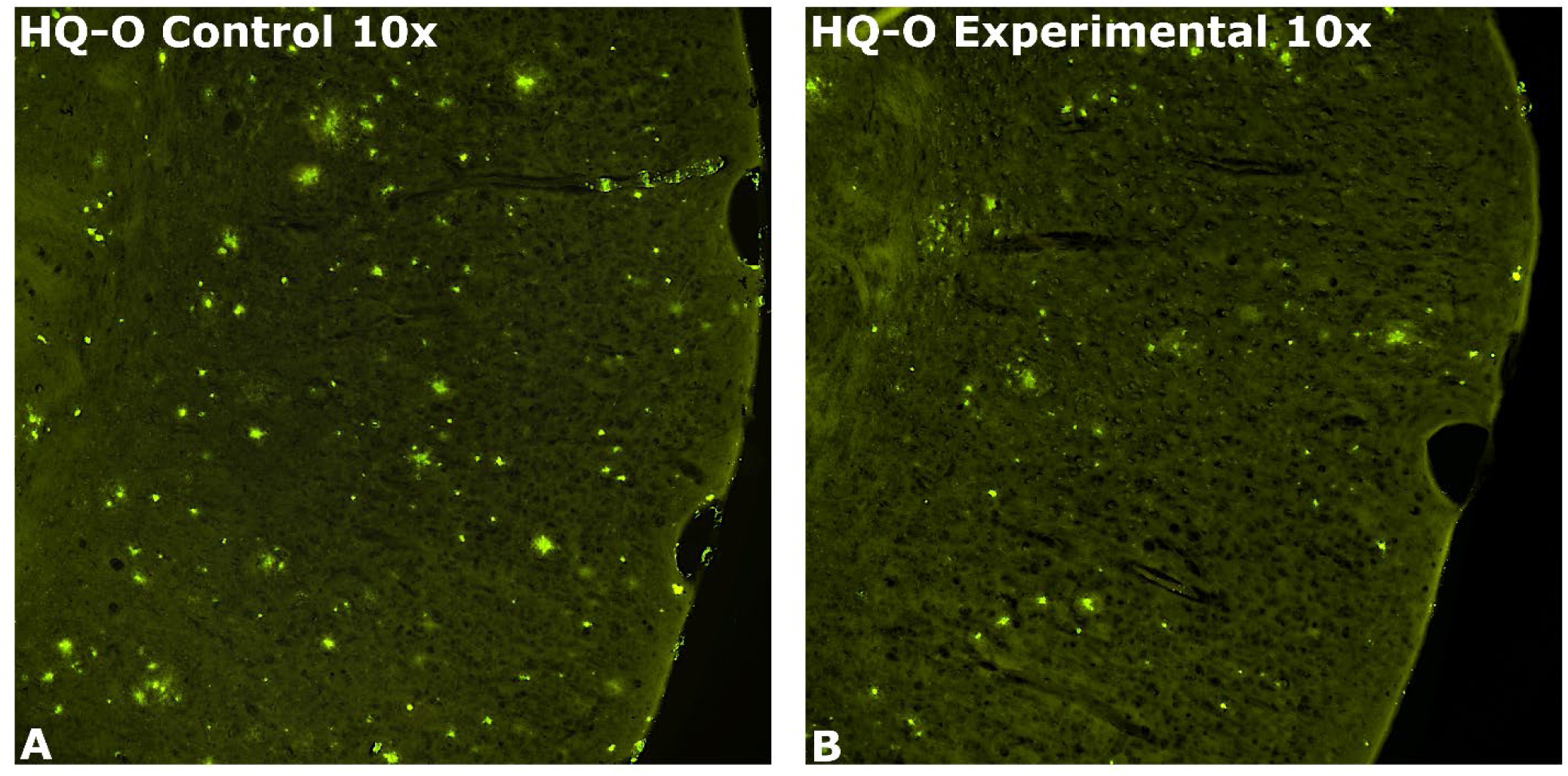
Typical HQ-O labeled amyloid plaques. A) Untreated control animals exhibit a high density of amyloid plaques in the temporal cortex. B) PAA administered animals show a significantly reduced amyloid plaque burden in the parietal cortex.

The combined plaque area in the control group was 41,312,162 µm², whereas the PAA-treated group exhibited a total plaque area of 27,931,580 µm², representing approximately 67% of the plaque burden observed in untreated mice (Fig. 3). Quantitative analysis demonstrated that PAA treatment significantly reduced total plaque area (Fig. 3A), plaque number (Fig. 3B), and mean plaque area (Fig. 3C) (all p ≤ 0.05). These findings indicate that PAA treatment reduced both the incidence of new plaque seeding and the growth of existing plaques.

**Fig. 3.**
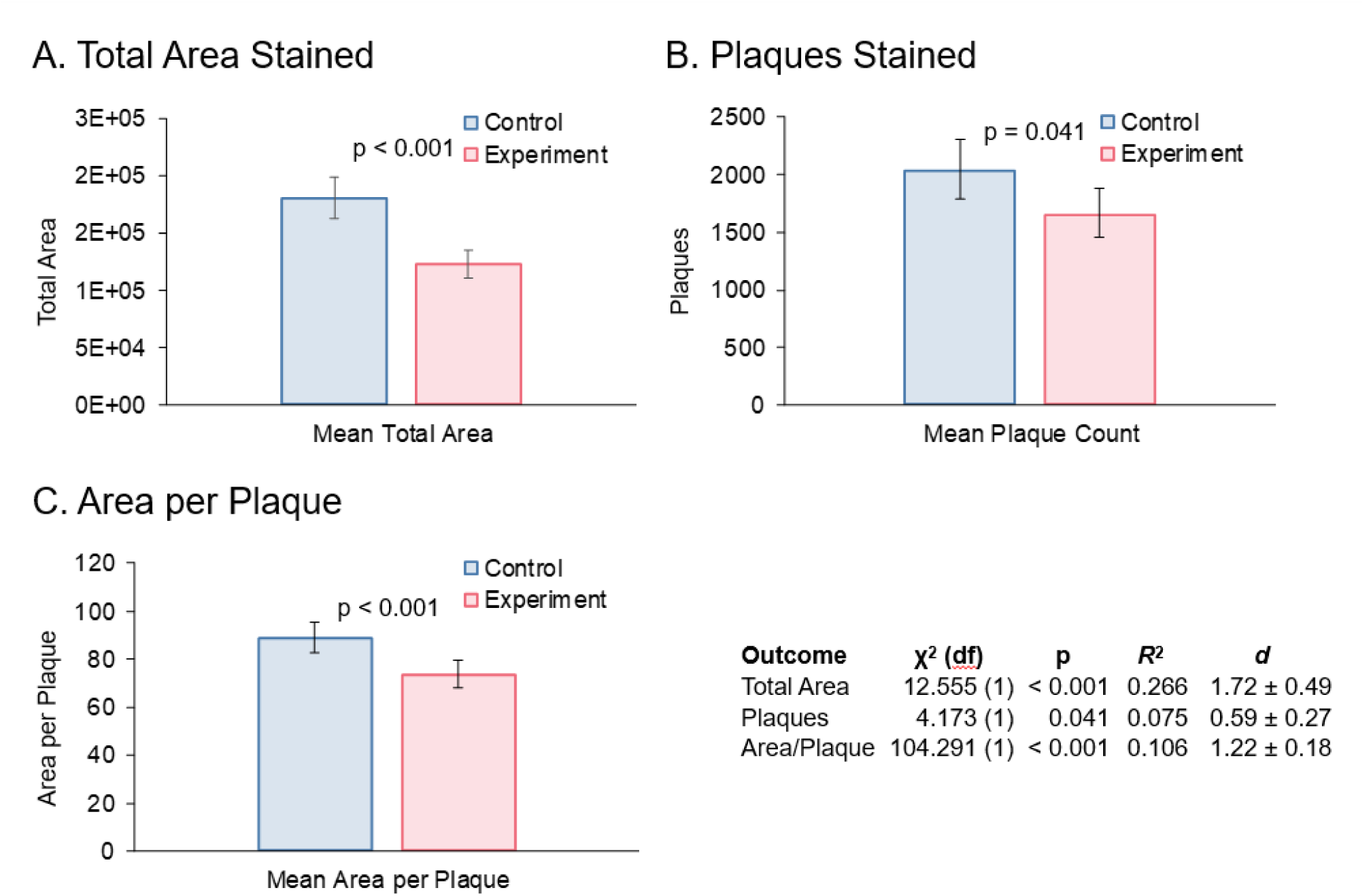
Effect of treatment of Aβ plaque in mouse brains. Mice were treated with PAA, brains harvested and imaged, as described herein. A) Marginal means and estimated standard errors are shown. Total area of Aβ plaques. B) Plaque counts. C) Mean area per plaque. Model ANOVAs, *R*^2^, and Cohen’s *d* are presented.

### Differential brain tissue labeling suggests that PAA recognizes distinct plaque-associated components

Incubation of brain tissue sections with PAA produced high-contrast, high-resolution red fluorescent labeling of all parenchymal amyloid plaques. PAA fluorescence was readily detected using both blue (Fig. 4A) and green (Fig. 4B) excitation. Post-incubation of the stained sections in EuCl₃ further enhanced fluorescence intensity under UV excitation (Fig. 4C).

**Fig. 4.**
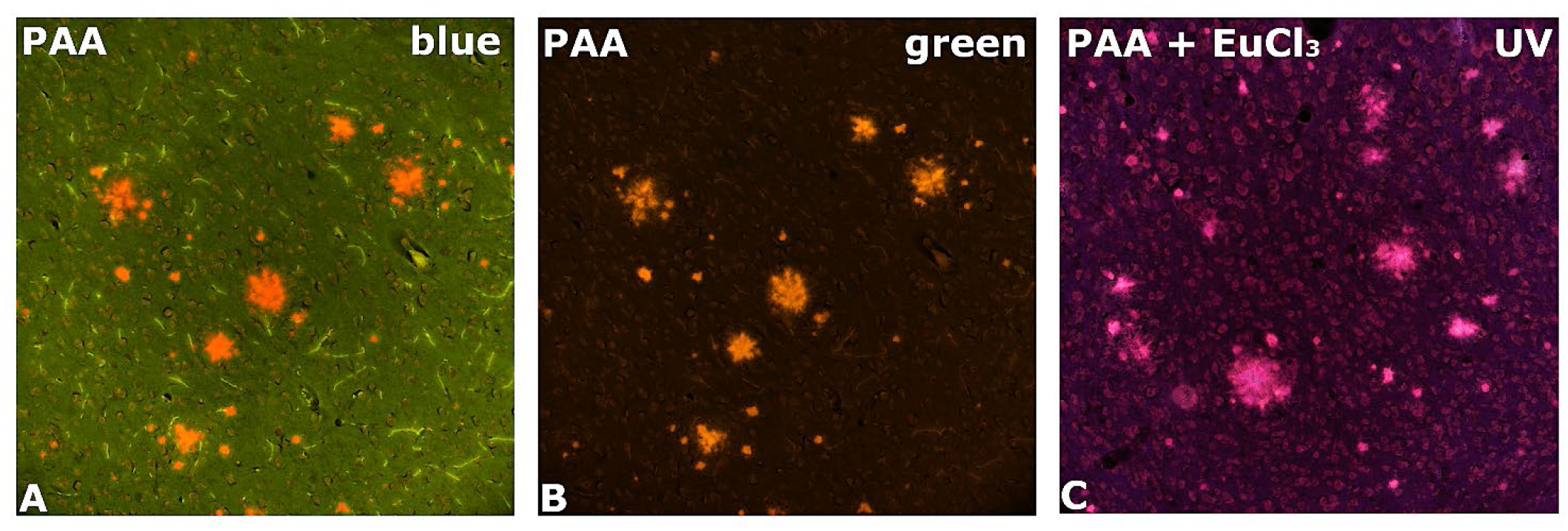
PAA staining of amyloid plaques. Fluorescent red stain under A) blue or B) green light excitation. C) Tissue sections incubated in solutions of PAA followed by EuCl_3_ show a red/pink staining of amyloid plaques under UV excitation.

Although PAA labeled all parenchymal plaques, it did not label vascular amyloid deposits. The labeled parenchymal plaques displayed a characteristic morphology consisting of a dense central core, a fibrillar intermediate region, and a more diffuse and globular peripheral region.

To compare plaque labeling patterns, PAA staining was combined with HQ-O labeling on the same tissue sections. In solvent-delipidized sections, the staining patterns of PAA and HQ-O were nearly identical (Fig. 5A–C). In contrast, in non-delipidized frozen sections, HQ-O labeling was largely confined to the central core and intermediate regions of the plaques, whereas PAA labeled not only these regions but also the surrounding peripheral zone (Fig. 5D–F). This broader labeling pattern suggests that PAA recognizes plaque-associated components that are not detected by HQ-O in intact frozen tissue.

**Fig. 5.**
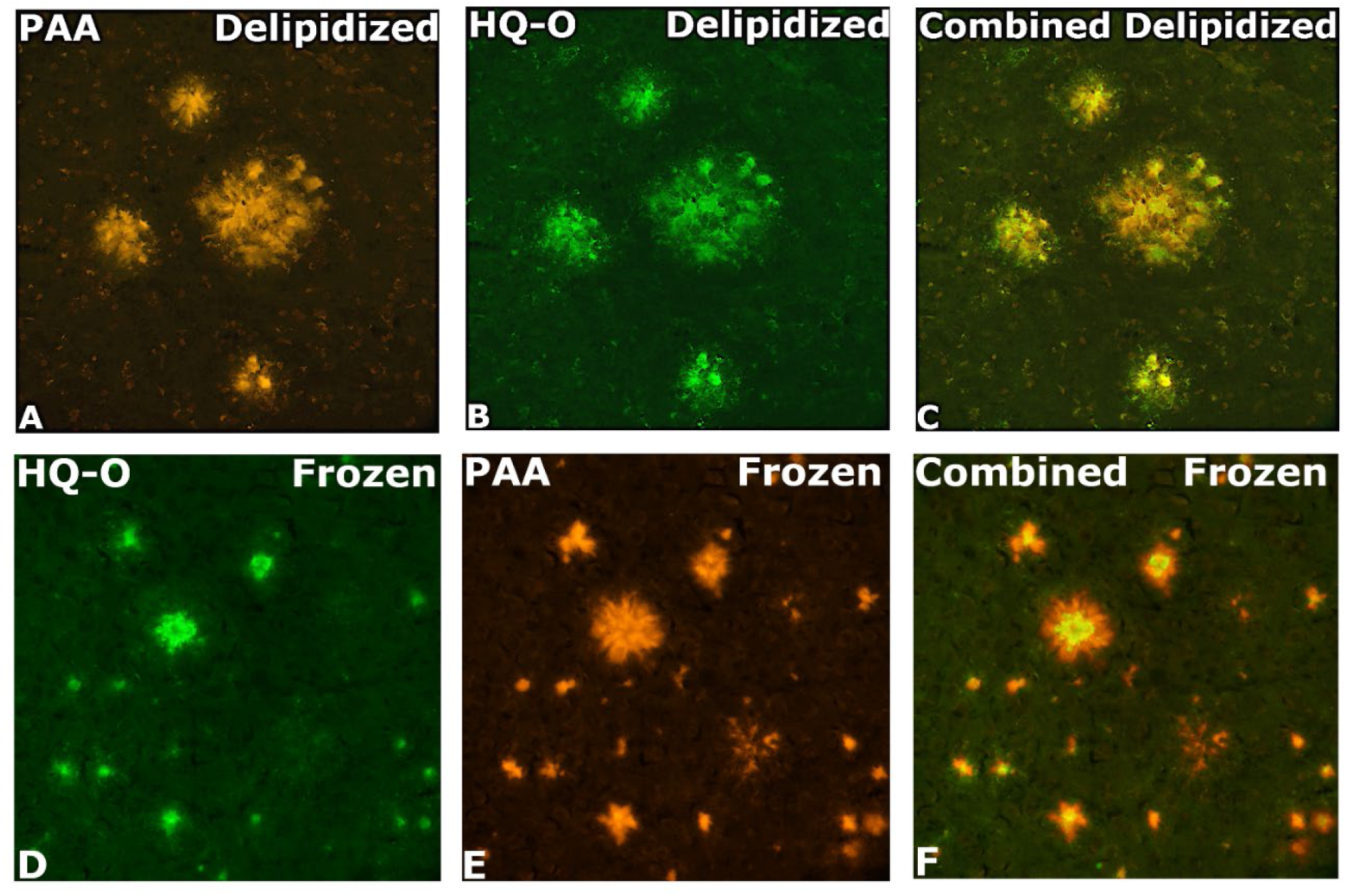
Combined PAA and HQ-O. Combined incubation of solvent delipidized tissue sections shows green appearing HQ-O labeled plaques under A) blue light excitation and B) red fluorescent plaque labeling is seen under green light excitation. Combining these images shows similarities and differences between the C) HQ-O and PAA staining patterns. Combined PAA and HQ-O incubation of frozen cut tissue sections shows green appearing HQ-O labeled plaques under D) blue light excitation. E) Red fluorescent plaque labeling under green light excitation. F) Combining these images shows similarities and differences between the HQ-O and PAA staining patterns.

## Discussion

This study demonstrates the effectiveness of oral dosing with PAA in reducing the amyloid plaque burden in mice models of AD. Unlike the previous study (71) which used an APP/Tau model of AD, this study used the APP/PS1 mouse model of AD to avoid the mortality associated with littermate aggression seen with the APP/Tau strain and because the APP/PS1 strain results in a substantially greater plaque burden, which is the primary endpoint of this study. Despite the different animal models used, the total average plaque reduction seen in the dosed vs. the untreated control animals was remarkably similar, confirming that PAA administration resulted in approximately 2/3 the total plaque areas compared to the untreated control animals in 2 different mouse models of AD.

The mechanism whereby daily oral administration of PAA, or certain other metal chelators, reduces the amyloid plaque burden in transgenic mouse models of AD has not been resolved definitively. Presumably the mechanism of action of PAA is based on its chemical properties, which include metal chelator, aromatic cation, and metalloprotease inhibitor (83). We proposed (71) an ability for PAA to chelate oligo-valent metals and thereby inhibit metal coordinated Aβ aggregation, or ‘metal seeding’ (Fig. 6). Transition metals with oxidation states of more than one, such as Fe, Zn, and Cu, have received the most attention as a potential nucleus for the aggregation of Aβ. It’s worth noting that PAA can form chelates with all 3 of these transition metals but is not expected to bind to alkali metals such as lithium, which has been shown to increase neuronal cell viability (84, 85).

**Fig. 6.**
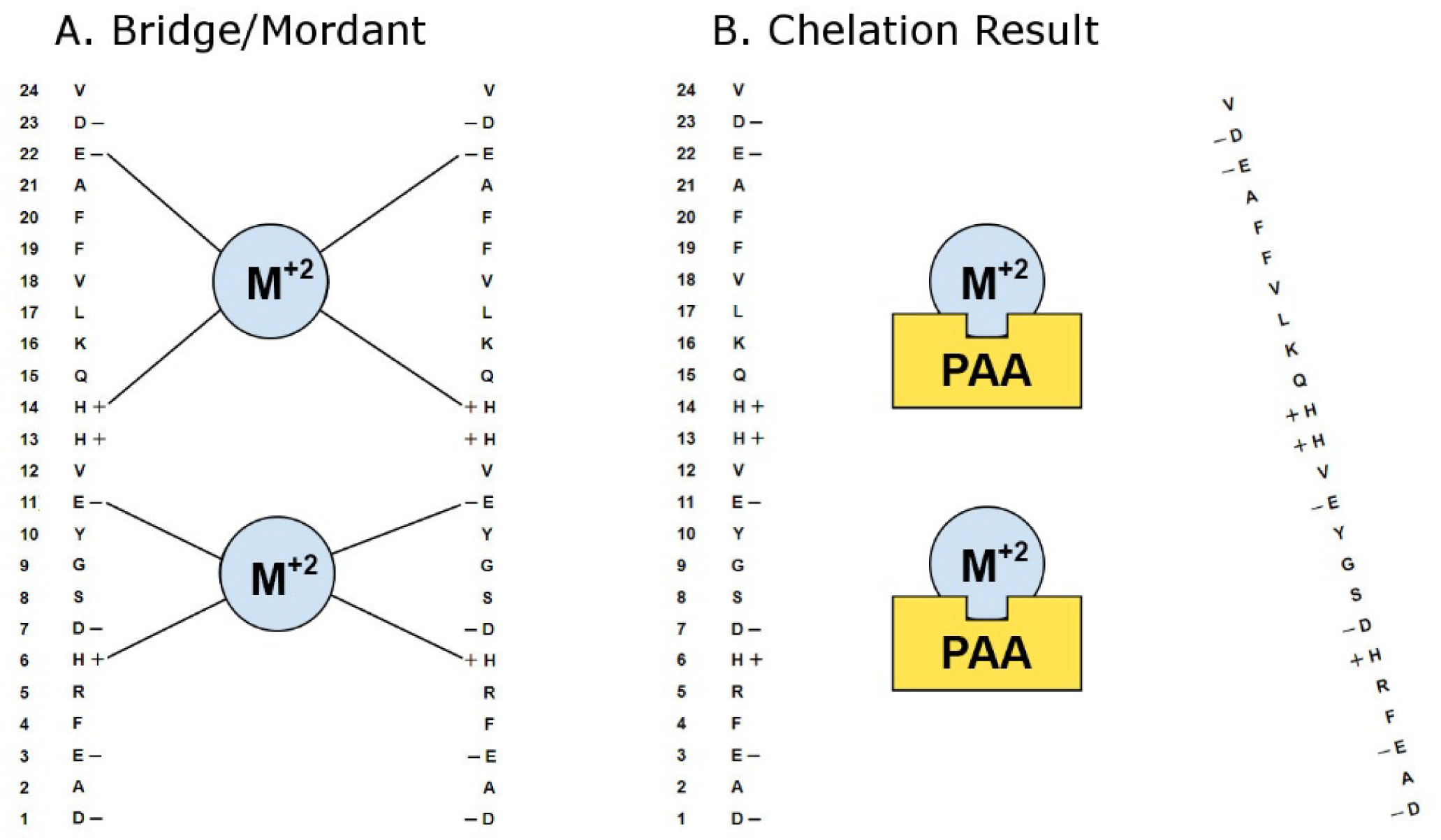
Seeding and interference by PAA. A.) Possible mechanism whereby certain transition metals (M^+2^), such as Fe or Zn, serve to coordinate or seed the aggregation of Aβ in AD. B.) Possible mechanism for the prevention of this metal induced aggregation by the metal chelator PAA. The columns of numbers and letters refer to the amino acid and its position within the Aβ peptide.

PAA differs from other metal chelators that have shown some promise in reducing the amyloid plaque burdens in mouse models of AD by not being amphoteric, as it contains only positively charged primary and tertiary nitrogen groups (Fig 1). It is conceivable that the absence of negatively charged carboxy or hydroxy groups facilitates blood-brain barrier (BBB) penetration. This would be consistent with the observation that many small positively charged dyes and drugs can cross the BBB, while most acidic dyes and drugs do not cross the BBB.

The cationic nature of PAA could underlie a second possible mode of action. Specifically, the positively charged amino group could bind to the negatively charged sialic acid moiety of glycolipids (Fig 7). It has been proposed Aβ binds to membrane bound gangliosides forming GAβs which undergo a conformational change from the α-helix to the *B*-sheet configuration (62, 86).

**Fig 7.**
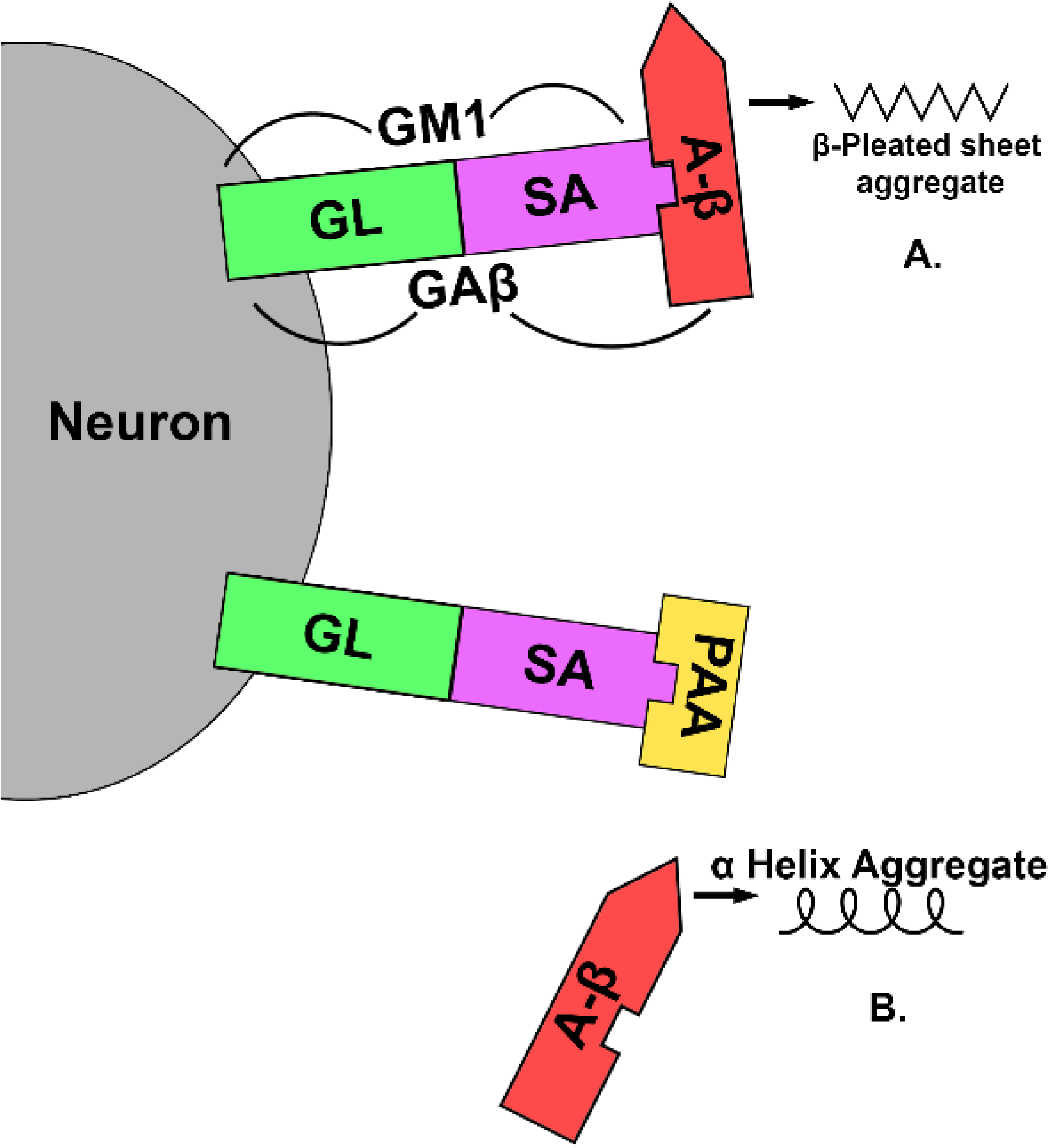
GAβ formation and interferences by PAA. A.) Possible mechanism whereby Aβ binds to the ganglioside GM1 to form GAβs which favor a β-pleated sheet configuration. B.) Inhibition of GAβ formation by PAA binding. GL = glycolipid; SA = sialic acid.

It is plausible that the positively charged amino group of the PAA molecule binds to the negatively charged carboxylic group of the sialic acid residues of the GM1 molecule, thus inhibiting the formation amyloidogenic GAβs. In such a scenario, the positively charged lysine within Aβ could bind to the sialic acid in GM1 to form GAβs, which might be prevented by the positively charged PAA binding to the sialic acid within membrane bound GM1 molecules. The notion that PAA binds to glycosylated molecules is supported by the observation that the PAA staining pattern is similar to the plaque labeling pattern seen when using probes for localizing glycosylated compounds such as the PAS procedure, the Ruthenium Red stain, and WGA lectin histochemistry, which all label molecules containing n-acetylneuraminic acid.

The two possible mechanisms of action discussed need not be mutually exclusive. For example, it is conceivable that, by virtue of its free amino group, PAA binds to the sialic acid component of membrane bound GM1 and could then chelate multivalent metals via its 2 tertiary nitrogen atoms, thereby inhibiting the metal coordinated aggregation of Aβ.

Our labeling studies using PAA combined with HQ-O revealed similarities and differences in their labeling patterns in frozen sections. Although both tracers labeled all parenchymal plaques, there were noticeable differences in their respective labeling patterns with the most conspicuous difference being that in addition to PAA staining everything that HQ-O stained, it also labeled a more peripheral portion of the plaques in frozen tissue sections. The reason for this differential staining is not yet known and might be interpreted as simple diffusion of the tracer from its target, but this seems unlikely because the staining pattern was stable and did not change as a result of variations in incubation or rinse times. It is therefore possible that this staining pattern signifies its binding to a non-fibular portion of the plaque, possibly glycolipids. This hypothesis is supported by the observation that lipid extraction, either by immersion of frozen sections in xylene and ethanol or via paraffin embedding, prevented the labeling of the more peripheral portions of the plaques seen in frozen tissue sections. This might also be the reason that PAA, like PAS and Ruthenium Red, stains parenchymal plaques but not vascular plaques.

Although the PAA staining/labeling described here was developed primarily to evaluate the affinity PAA has for amyloid plaques, it may also have general histological applications for the simple, rapid, high resolution, and high contrast staining of amyloid plaques. Potential advantages include labeling a greater portion of the plaques in frozen tissue sections than seen with probes targeting Aβ aggregates. Also, the resulting red fluorescence is much brighter than other red amyloid binding fluorochromes such as Congo Red.

A number of follow-up studies could further clarify the potential role of PAA in the study and treatment of AD. One valuable follow-up study could involve looking for behavioral correlates. Other useful follow-up studies could include toxicological and pharmacokinetic endpoints. It would also be interesting to examine phenanthroline derivatives with alternative functional groups for possible amyloid-inhibiting properties, which could help resolve the underlying target and mode of action of the compound. To help resolve PAA’s mode of action tissue culture-based studies are ongoing. Preliminary results showed PAA did not adversely affect cell viability under the experimental conditions (data not shown). CellTiter-Glo (CTG) viability and lactate dehydrogenase (LDH) cytotoxicity assays demonstrated that PAA was well tolerated at concentrations up to 3 nM in differentiated neuronal cultures (data not shown). We noted that PAA altered the pH and taste of the water. Consumption did not differ between treatment groups, indicating no gross avoidance. However, it may have been appropriate to have had pH matched (HCl-treated) water for the control group, particularly given that stronger HCl acidification (pH 2.8) produced significant behavioral and neuropathology improvements in a non-AD mouse model. A three-level treatment of water/acidified water/PAA may resolve this, admittedly very fine-grained, question (87).

In conclusion, this study verified that oral dosing of APP/PS1 mice with PAA significantly reduced the area of amyloid plaques compared to untreated control mice. It also provides support for the theory that PAA has the potential to bind to metals such as Fe and Zn intercalated within Aβ aggregates and also has the potential to bind to sialic acid in plaque-bound gangliosides. Thus, PAA could be of value both as a potential AD therapeutic agent and as a fluorescent histological probe for localizing amyloid plaques in tissue sections.

## Conclusions

We confirmed that oral dosing with PAA resulted in plaque reductions comparable to that seen in the original study. We further demonstrated that PAA can bind directly to amyloid plaques. PAA, as an all-cationic metal chelator, could be binding to either transition metals incorporated within the amyloid plaques or to the sialic acid portion of gangliosides.

## List of Abbreviations

AD: Alzheimer’s disease;
Aβ: amyloid beta;
BBB: blood-brain barrier;
GM1: monosialotetrahexosylganglioside;
HQ-O: 8-hydroxyquinoline oxalate;
PAA: 1,10-phenanthroline-5-amine;
PAS: periodic acid Schiff’s-reagent;
SMON: subacute myelopathic neuropathy.

## Acknowledgements

The authors thank partial support from NIH and Indiana University Distinguished Professor fund.

## Authors Contributions

LS: Conceptualization, Methodology, Validation, Investigation, Resources, Writing, Visualization, Supervision, Project administration, Funding acquisition; BM: Formal Analysis, Visualization, Writing; CS: Methodology, Validation, Investigation, Resources, Visualization; KG: Tissue culture experiments, DKL: Planning, Writing, Revisions, and Submission.

## Statements and declarations

### Ethical considerations

No human participants. All animal experimental procedures were conducted in strict accordance with the Guide for the Care and Use of Laboratory Animals and were approved.

### Consent to participate

Not applicable.

### Consent for publication

Not applicable.

## Availability of Data and Materials

The data supporting the findings of this study are available on request from the corresponding author. The data is not publicly available due to privacy or ethical restrictions.

